# RHO GTPase of Plants contributes to robust establishment of cellular asymmetry during development from a single cell

**DOI:** 10.64898/2026.08.18.745417

**Authors:** Hugh Mulvey, Yuuki Sakai, Katharina Jandrasits, Zohar Meir, Magdalena Mosiolek, Kimitsune Ishizaki, Liam Dolan

## Abstract

A fundamental question in developmental biology is how highly complex, yet reproducible multicellular body plans form from a single cell. The multicellular haploid body of the land plant *Marchantia polymorpha* develops from a single isolated cell – the spore – that divides asymmetrically. This produces a small terminally differentiating basal cell, and a large proliferative apical cell that gives rise to the multicellular sporeling body. The genetic basis for the establishment of asymmetry, first within the spore, and later within the sporeling body has remained virtually unknown. Here, we show that the plant specific RHO-type GTPase – RHO OF PLANTS (ROP) – polarises to the spore basal pole to ensure spore division is highly and reproducibly asymmetric. We demonstrate this highly asymmetric division is necessary to specify the terminal differentiation of the basal cell. Furthermore, we show that ROP-mediated polarised outgrowth is required to establish asymmetry within the sporeling body derived from the apical cell. Our discovery highlights how ROP function confers developmental robustness and contributes to the establishment of asymmetry during plant development from a single isolated cell.

## INTRODUCTION

The morphogenesis of a complex multicellular body from a single cell requires the generation of diverse cell types, spatially arranged into tissues and organs. Asymmetric cell division is often correlated with the generation of different cell types^1,2^. In the model organisms *Caenorhabditis elegans* and *Arabidopsis thaliana*, the single cell which gives rise to the multicellular diploid body – the zygote – divides asymmetrically to form cells with different developmental fates^3,4^. In both cases, the zygote develops surrounded by parental tissue, and cell polarisation, which precedes asymmetric cell division, is influenced by parental factors^3,4^. The complex thalloid liverwort *Marchantia polymorpha* (Marchantia), also initiates development with an asymmetric cell division at the one cell stage. However, unlike *C. elegans* or *A. thaliana*, this first cell – the haploid spore – develops in complete isolation from parental cells and tissues. Marchantia therefore offers a unique opportunity to understand how cell polarity and developmental asymmetry emerge in the absence of parental cues.

Marchantia spores are produced in tetrads when the diploid spore mother cell undergoes meiosis^5^. Within the tetrad, the four spores are tetrahedral in shape^6^. However, after they separate into individual spores and mature, they become spherical and lack any obvious signs of morphological asymmetry^6^. The spores then grow isotropically, expanding in diameter from around 10 µm to 20–30 µm, before dividing asymmetrically to produce a small basal cell and a large apical cell (Figures 1A and 1B). These two cells have contrasting fates: the basal cell usually undergoes tip growth to terminally differentiate into the first rhizoid cell (germ rhizoid), whilst the apical cell continues to divide to give rise to the early cell mass^7^. The early cell mass develops into an asymmetric structure, and eventually the prothallus – the flat plant body – develops at its apex^8^. The genetic basis for the establishment of asymmetry initially within the spore, as well as later within the early cell mass, remains unknown. Furthermore, the significance of cell size asymmetry for cell fate asymmetry following asymmetric spore division, has never been reported in Marchantia.

**Figure 1.**
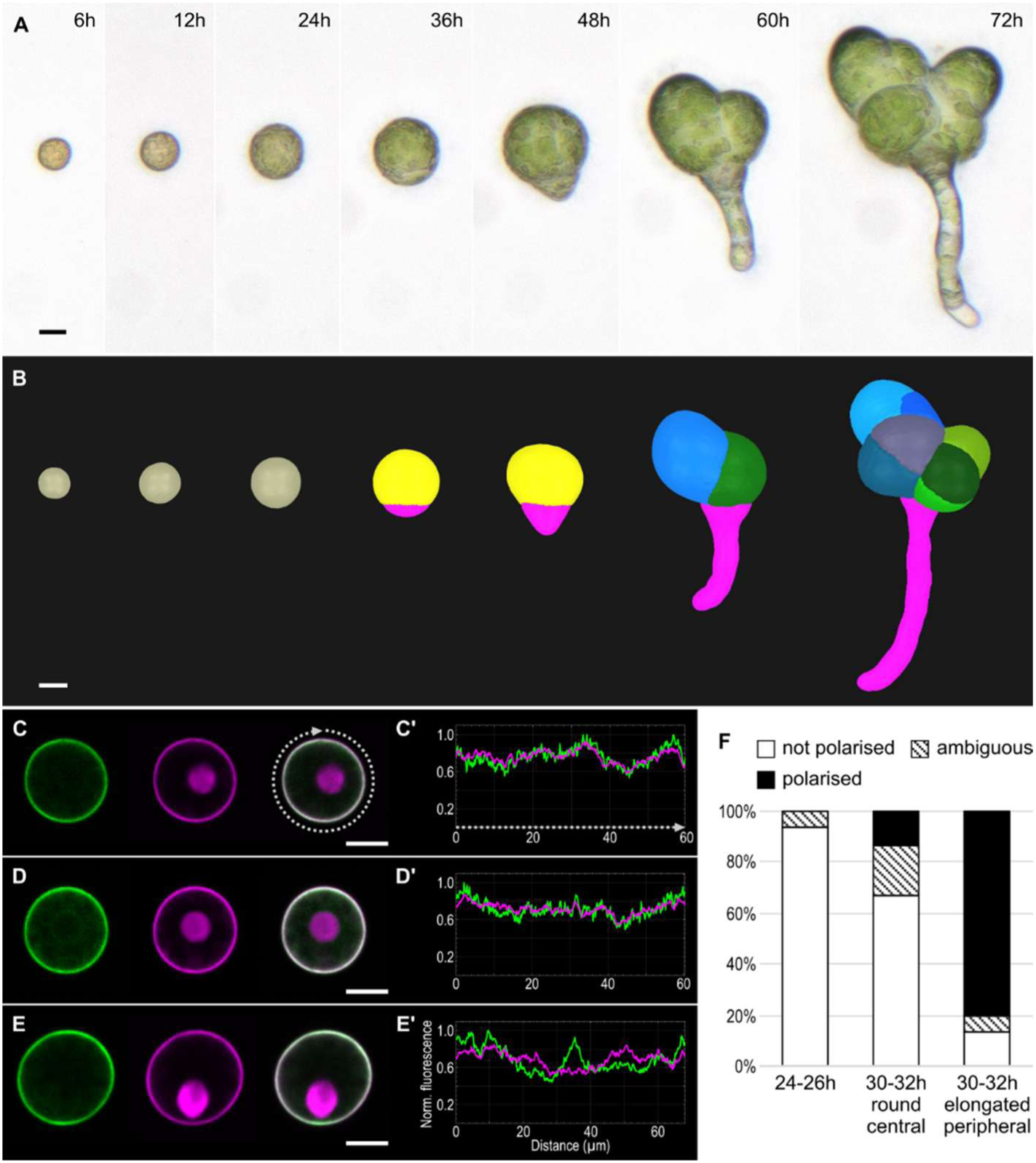
Venus-MpROP polarises to the basal pole before asymmetric spore division. **(A–B)** The *Marchantia polymorpha* spore grows isotropically then divides asymmetrically to give rise to two cells with distinct fates. (A) Time lapse images of an individual spore 6–72 h after plating on media. (B) 3D segmentation of spores and sporelings fixed at the time points shown above in A. **(C–E)** Representative images of spores expressing *_pro_*Mp*ROP:Venus-*Mp*ROP*, *_pro_*Mp*UBE2:mScarletI-*At*LTI6b*, and *_pro_*Mp*ROP:mScarletI-N7* from (C) 24–26 h, (D) 30–32 h with a round central nucleus, (E) 30–32 h with an elongated peripheral nucleus. From left to right: Venus, mScarletI, and merged fluorescence images. **(C’–E’)** Normalised fluorescence intensity profiles of Venus-MpROP (green) and mScarletI-AtLTI6b (magenta) along the plasma membrane (clockwise from 12 o’clock as indicated by the grey dotted lines in C and C’) for the spores in C–E. **(F)** Proportions of spores in which polarised Venus-MpROP was detected. n = 15 for each of the three categories. Scale bars, 10 µm in (A–E).

The asymmetry of spore division implies that the spore is polarised before it divides. Consistently, the microtubule and actin cytoskeleton are polar localised within the spore and promote the migration of the nucleus to the basal pole before asymmetric spore division^9,10^. However, what causes the cytoskeleton to polarise in the first place remains unclear. Phototropin-mediated blue-light signalling orients the spore polarity axis, with the basal pole forming on the shaded side of the spore that is illuminated with unidirectional blue light^11^. In the absence of Phototropin function, spore divisions are randomly oriented but remain asymmetric. This suggests that blue light acts as a spatial cue that aligns the apical-basal polarity axis to the external environment, but is not required to establish polarity. Whether some other spatial cue initially polarises the spore remains unknown, raising the possibility that the Marchantia spore can polarise *de novo*.

The RHO family of GTPases regulates cell polarity and cytoskeletal organisation in diverse eukaryotes^12^. For example, in the budding yeast *Saccharomyces cerevisiae*, the RHO GTPase CDC42 plays an essential role in polarising the cell and promoting actin-mediated bud outgrowth^13,14^. Through a positive feedback mechanism involving one of its core activators, CDC42 polarises to a single cortical site marked by landmark proteins inherited from the mother cell^15^. Even in the absence of landmark proteins or other spatial cues, this feedback mechanism allows CDC42 to polarise spontaneously to a random cortical site, consistent with the Turing model of *de novo* pattern formation^16,17^. Although the plant specific RHO subfamily – RHO OF PLANTS (ROP) – has diverged from its counterparts in fungi and animals, the core RHO signalling module with its regulators, are conserved in land plants including Marchantia^18,19^. This led us to hypothesise that ROP may function in polarising the Marchantia spore.

Uniquely among land plants, ROP and its core regulators are encoded by single copy genes in Marchantia^20,21^. Furthermore, ROP signalling regulates organogenesis during the vegetative phase of Marchantia development^22–24^. We therefore hypothesised that ROP functions not only in establishing spore polarity, but also in polarising the subsequent development of the early cell mass.

Here, we report our discovery that ROP polarity contributes to the robustness of asymmetric spore division. Furthermore, utilising Mp*rop* mutant spores, we demonstrate the requirement of spore division asymmetry for cell fate asymmetry. Finally, we show that ROP contributes to the establishment of asymmetry within the sporeling early cell mass.

## RESULTS

### MpROP polarises to the basal pole before asymmetric spore division

If the MpROP protein functions in polarising the spore, we hypothesised that it would be enriched at the cell cortex, at one side of the cell, before asymmetric spore division. To assess if the MpROP protein is polar localised at the cell cortex, we imaged spores expressing Venus-MpROP and mScarletI-AtLTI6b (a uniform plasma membrane marker) and compared the Venus and mScarletI signal intensity profiles along the cell perimeter (Figures 1C–1E’). At 24–26 h after plating on solid media (4–6 h before most spores enter division), the spores appeared round in cross section, and a single round nucleus (in most cases centrally positioned) was visible (Figure 1C). At this stage, Venus and mScarletI signal intensity profiles overlapped and no specific enrichment of Venus signal was detected along the cell perimeter, indicating that Venus-MpROP was localised uniformly at the cell cortex (Figures 1C’ and 1F).

At 30–32 h, in spores with a round centrally positioned nucleus, polarisation of Venus-MpROP was detected in a minority of cases (Figure 1F). By contrast in spores with an elongated nucleus positioned at the cell periphery, indicative of the basal migration of the nucleus, Venus-MpROP was enriched at the basal pole in the majority of cases (Figures 1E, 1E’, and 1F). These results indicate that MpROP is polarised to the basal pole before mitosis and is therefore consistent with the hypothesis that MpROP functions in spore polarisation.

### MpROP ensures robust asymmetric cell division of the spore

Since Venus-MpROP is polarised to the basal pole before mitosis, we hypothesised that MpROP contributes to the development of division asymmetry. To determine if spore polarisation is defective in the absence of MpROP function, we attempted to generate an isogenic population of Mp*rop* loss of function mutant spores by crossing male and female plants carrying the same complete loss-of-function mutation (Mp*rop-3*). This cross failed, likely due to the severe morphological defects of the Mp*rop-3* mutants that led to sterility^22^. We therefore generated a segregating population of Mp*rop-3* and wild-type spores by crossing a male Mp*rop-3* with a female wild type (Tak-2) (Figure S1A).

We first tested the viability of Mp*rop-3* spores produced from the cross between Mp*rop-3* and Tak-2. The germination rate of spores produced from Mp*rop-3* x Tak-2 and Tak-1 x Tak-2 was similar, suggesting that Mp*rop-3* spores are as viable as wild-type spores (Figures S1B–S1E).

Furthermore, one week after plating spores from the Mp*rop-3* x Tak-2 cross, 47% developed very short or no visible rhizoids – a distinctive phenotype of Mp*rop-3* gametophyte^22^ – consistent with the expected 1:1 segregation of Mp*rop-3* and wild-type progeny (Figures S1C–S1E). Genotyping confirmed sporelings with very short or no rhizoids as Mp*rop-3* and those with long rhizoids as wild type (Figure S1F). Thus, the cross between Mp*rop-3* and Tak-2 provided a viable segregating spore population to assess the effect of the loss of MpROP function on spore division asymmetry.

To determine if MpROP function is required for the development of spore asymmetry, we quantified spore division asymmetry as the volume of the small (basal) cell relative to the total volume of a recently divided spore (relative basal cell volume). In the pure wild-type population, the majority of spore divisions were highly asymmetric and symmetric division (defined as those resulting in a relative basal cell volume > 35%) was observed in less than 3% of spores (Figures 2A, 2B, and 2E). In the Mp*rop-3*-wild-type segregating population, the majority of spore divisions were also asymmetric, however, the average relative basal cell volume was significantly greater than in the pure wild-type population, indicating a tendency for spores in the segregating population to divide more symmetrically (Figures 2A, 2C, and 2E). Furthermore, symmetric divisions occurred at a higher frequency in the segregating population (8.6%) than in the pure wild-type population (2.9%) (Figure 2E). These results are consistent with the hypothesis that the loss of MpROP function causes a reduction in spore division asymmetry.

**Figure 2.**
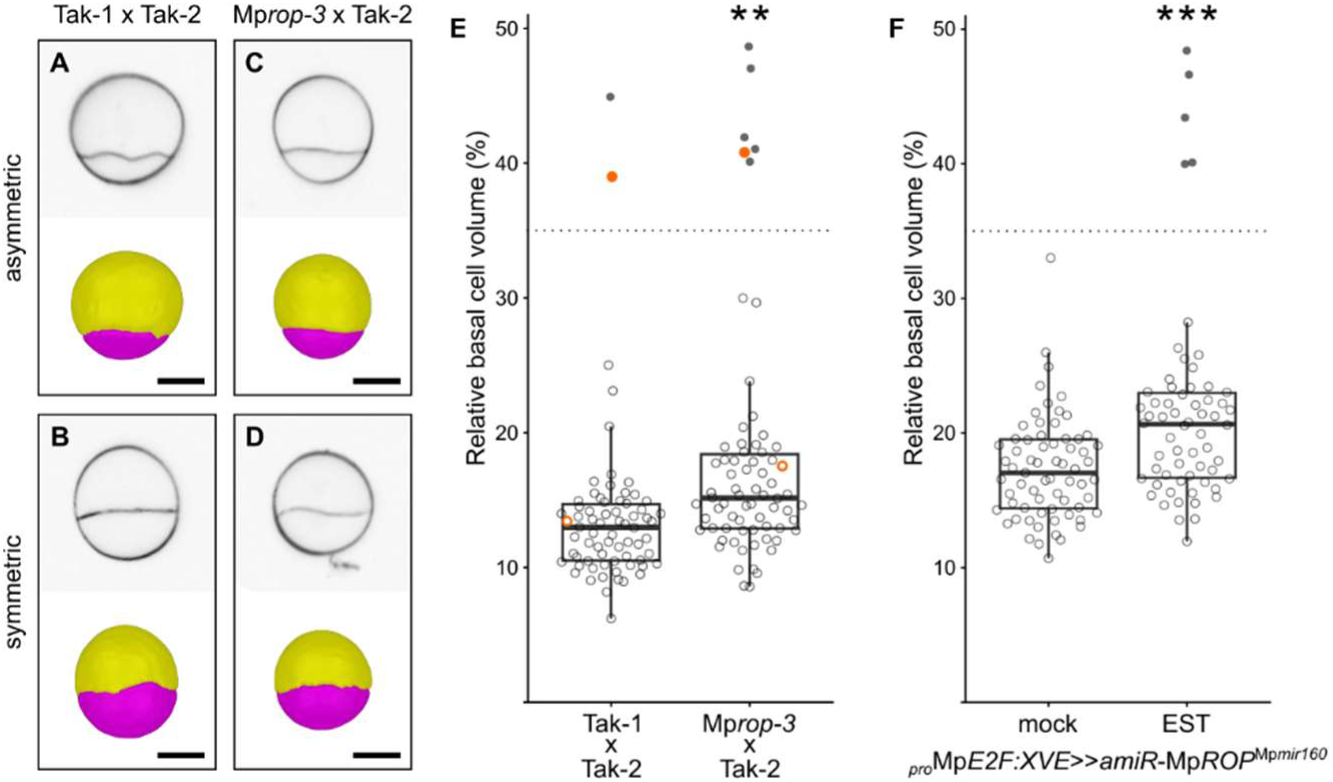
MpROP function ensures robust asymmetric cell division of the spore. **(A–D)** Representative asymmetrically and symmetrically divided spores from the pure wild-type population (Tak-1 x Tak-2) and the segregating Mp*rop* mutant*-*wild-type population (Mp*rop-3* x Tak-2). Upper panels: Optical median sections along the apical-basal axis. The cell wall (black) was stained with the SR2200 dye after sample fixation and clearing; Lower panels: 3D segmentation of samples in the upper panels. Larger (apical) cell in yellow and smaller (basal) cell in pink. Scale bars, 10 µm. **(E)** Relative basal cell volume (basal cell volume/total volume) in recently divided spores (36 or 40 h after plating) was significantly greater in the Mp*rop-3* x Tak-2 population than in the Tak-1 x Tak-2 population (Welch’s t-test, p = 0.0016; n = 70 for each population). Filled circles indicate individuals classified to have divided symmetrically (relative basal cell volume > 35%, dotted line). Symmetric division was more frequent in Mp*rop-3* x Tak-2 (8.6%) than in Tak-1 x Tak-2 (2.9%), although the difference was not significant (p = 0.137, one-sided Fisher’s exact test). Data points for samples shown in A–D are marked in orange. **(F)** Relative basal cell volume in recently divided spores (40 h after plating) expressing *_pro_*Mp*E2F:XVE>>amiR-*Mp*ROP*^Mp*mir160*^ was significantly greater following estradiol (EST) treatment than mock (DMSO) treatment (Welch’s t-test, p = 1.7 × 10⁻^4^; n = 67 mock, n = 62 EST). Frequency of symmetric division was significantly higher with EST (8.1%) than without (0%) (p = 0.024, one-sided Fisher’s exact test). **p < 0.01; ***p < 0.001 for Welch’s t-test. See also Figures S1 and S2.

To independently assess the involvement of MpROP function in asymmetric spore division, we generated an isogenic population of inducible Mp*ROP* knock-down spores (Figure S2). Spores expressing *_pro_*Mp*E2F:XVE*>>*amiR*-Mp*rop*^Mp*mir160*^ were treated either with estradiol to induce the expression of an artificial microRNA (amiR) that targets Mp*ROP* transcripts, or with DMSO as a mock treatment. The average relative basal cell volume was significantly greater in the estradiol-treated samples than in the DMSO-treated samples, consistent with the difference observed between Mp*rop-3* x Tak-2 and Tak-1 x Tak-2 populations (Figure 2F). Furthermore, the frequency of symmetric divisions was increased upon estradiol treatment (8.1%) compared to the DMSO control (0%) (Figure 2F). We therefore conclude that although not essential for asymmetric spore division, MpROP function ensures spore division is highly asymmetric and that this asymmetric division takes place in a robust and reproducible manner.

### MpROP promotes tip growth of the basal cell after asymmetric spore division

Since MpROP contributes to the development of cellular asymmetry, and by extension, to asymmetric spore division, we tested if MpROP specifies the distinct identities of the two daughter cells. Typically, in wild type, the apical cell divides and the basal cell undergoes tip growth to terminally differentiate into the germ rhizoid cell without further division (Figures 1A and 1B). We hypothesised that Venus-MpROP would remain polarised at the basal pole after asymmetric spore division to promote basal cell tip growth. Time-lapse imaging revealed that while Venus-MpROP was enriched at the basal pole before spore division, this enrichment decreased immediately after division (Figures 3C and 3D; n = 8/11 cases). During cytokinesis, Venus-MpROP marked the expanding cell plate and its polarisation at the basal pole was lost in most cases (Figure S3; n = 5/6). A few hours after spore division, Venus-MpROP became weakly polarised broadly along the basal cell surface (Figure 3F). Venus-MpROP polarisation became stronger and more confined to the basal pole in the following hours, as tip growth initiated at the basal pole (Figures 3G–3I). A change in sporeling shape associated with tip growth was detected only after Venus-MpROP polarisation (Figure 3J; n = 10/10). Taken together, these data indicate that MpROP is enriched at the basal pole before mitosis, loses this polar enrichment during cytokinesis, and then re-accumulates at the basal pole of the basal cell. This localisation at the pole of the basal cell suggests a causal link between Venus-MpROP polarisation and the initiation of basal cell tip growth.

**Figure 3.**
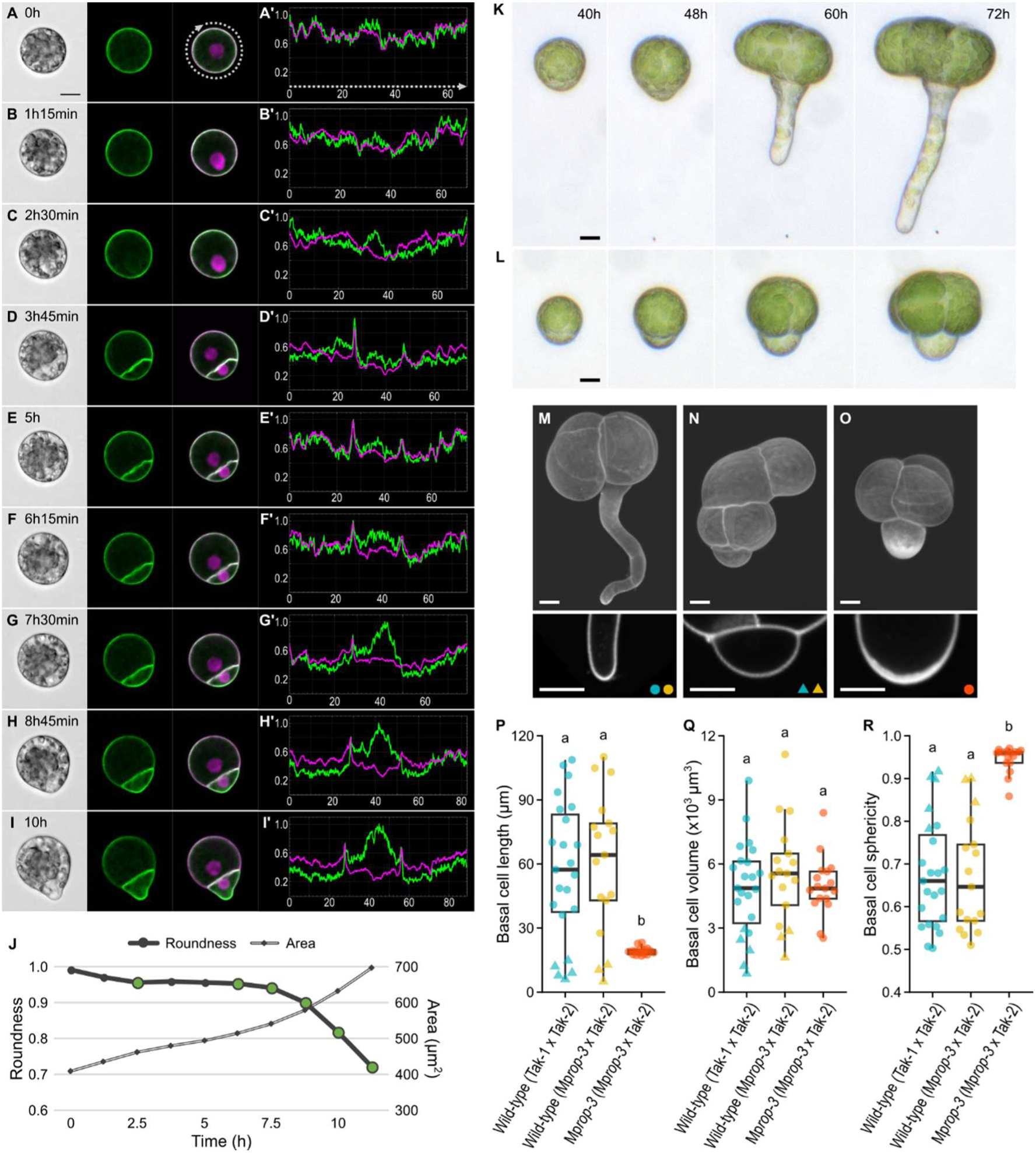
MpROP promotes tip growth of the basal cell following asymmetric spore division. **(A–I)** Time-lapse images of a representative spore expressing *_pro_*Mp*ROP:Venus-*Mp*ROP*, *_pro_*Mp*UBE2:mScarletI-*At*LTI6b*, and *_pro_*Mp*ROP:mScarletI-N7*. From left to right: brightfield, Venus, and merged fluorescence images. **(A’–I’)** Normalised fluorescence intensity profiles of Venus-MpROP (green) and mScarletI-AtLTI6b (magenta) along the spore or sporeling perimeter. Normalised fluorescence intensity on the y-axis and distance (µm) on the x-axis. **(J)** Change in spore/sporeling size (cross-sectional area) and shape (roundness) over time, measured from the brightfield images in A–I. Green dots mark time points when Venus-MpROP polarisation at the basal pole was detected. **(K)** Sporeling with tip growing basal cell, found in both Tak-1 x Tak-2 and Mp*rop-3* x Tak-2 populations. **(L)** Sporeling with isotropically growing basal cell, found only in the Mp*rop-3* x Tak-2 population. **(M–O)** Upper panels: 3D projections of fixed, cleared, and SR2200-stained 72-h-old sporelings. Lower panels: optical cross sections through basal cell tips. Blue and yellow symbols indicate basal cell morphologies observed in Tak-1 x Tak-2 and Mp*rop-3* x Tak-2 populations, respectively (M, N). Orange symbol indicates basal cell morphology observed only in the Mp*rop-3* x Tak-2 population (O). **(P–R)** Basal cell length (P; Welch’s ANOVA: p = 5.4 x 10^-7^), volume (Q; Welch’s ANOVA: p = 0.589); and sphericity (R; Welch’s ANOVA: p = 6.2 x 10^-12^), quantified for fixed and cleared 72-h-old sporelings. Mp*rop-3* x Tak-2 progeny were classified based on basal cell morphology: as wild type as in M (yellow circle) and N (yellow triangle), or Mp*rop-3* as in O (orange circle). Different letters indicate significantly different means (Games-Howell post hoc test, *p* < 0.05). n = 23 for wild type (Tak-1 × Tak-2), 17 for wild type (Mp*rop-3* × Tak-2), and 17 for Mp*rop-3* (Mp*rop-3* × Tak-2). All scale bars, 10 µm. See also Figures S3 and S4.

To test if MpROP function is required for basal cell tip growth, we tracked basal cell development following asymmetric spore division in populations derived from Tak-1 x Tak-2 and Mp*rop-3* x Tak-2 crosses. In the Tak-1 x Tak-2 population, tip growth initiated by 48 h and the basal cell elongated (aspect ratio > 2) by 60 h in most cases (Figures 3K and S4). Basal cell tip growth (or any noticeable basal cell growth) failed to initiate within the first 86 h in a minority of wild-type sporelings (Figure S4). In the asymmetrically divided Mp*rop-3* x Tak-2 population, the proportion of sporelings with an elongated basal cell was less than in the Tak-1 x Tak-2 population (Figure S4), and a subpopulation instead developed an isotropically-growing basal cell (Figure 3L). This is consistent with the hypothesis that MpROP function is required for basal cell tip growth.

To quantify the effect of the loss of MpROP function on basal cell shape, we imaged cell-wall-stained 72-h-old sporelings from the Tak-1 x Tak-2 and Mp*rop-3* x Tak-2 populations for 3D segmentation. Staining of the basal cell wall was mostly uniform in all Tak-1 x Tak-2 sporelings, regardless of whether tip growth had taken place (Figures 3M and 3N). By contrast, in the Mp*rop-3* x Tak-2 population, staining was highly enriched at the base of the basal cell specifically in sporelings where the basal cell had apparently grown isotropically (Figure 3O). Optical cross section revealed a thickened cell wall at the base of the basal cell in these sporelings. Given that this distinctive phenotype was observed in roughly 50% of sporelings in the Mp*rop-3* x Tak-2 population and never in the Tak1 x Tak-2 population, we inferred that these were Mp*rop-3* mutants. Based on this classification, Mp*rop-3* basal cells were significantly shorter than wild type basal cells (Figure 3P), but did not differ significantly in volume (Figure 3Q). Consequently, Mp*rop-3* basal cells had a higher degree of sphericity (Figure 3R). Thus, MpROP function is not required for the initial growth of the basal cell, but rather for polarising the growth of the basal cell.

### Highly asymmetric spore division, ensured by MpROP, promotes terminal differentiation of the basal cell

After establishing that MpROP promotes basal cell tip growth, we investigated the role of asymmetric spore division in specifying basal cell fate. To test if the physical asymmetry of spore division is required to specify the terminal differentiation of the basal cell into a rhizoid cell, we first compared the development of wild-type spores that had either divided asymmetrically or symmetrically.

The majority (97%) of spores from the Tak-1 x Tak-2 cross divided asymmetrically (Figure 2F). When we tracked the development of 59 asymmetrically divided wild-type spores, the larger apical cell alone continued to divide in most cases (n = 56/59; 95%), and both daughter cells divided in only three cases (5%) (Figures 4A, 4B, and 4G). By contrast, in 22 symmetrically divided wild-type spores, the division of both daughter cells was observed in 21 cases (95%) (Figures 4C and 4G). Furthermore, after a symmetric spore division, chloroplast content was similar in both daughter cells, and neither underwent tip growth (n = 22/22; 100%). This contrasts with asymmetric spore division, where the daughter cells with fewer chloroplasts – always the basal cell – underwent tip growth (n = 50/59; 85%). These data demonstrate that the generation of a smaller basal cell through asymmetric spore division is required to specify its terminal differentiation as a rhizoid cell without further division.

**Figure 4.**
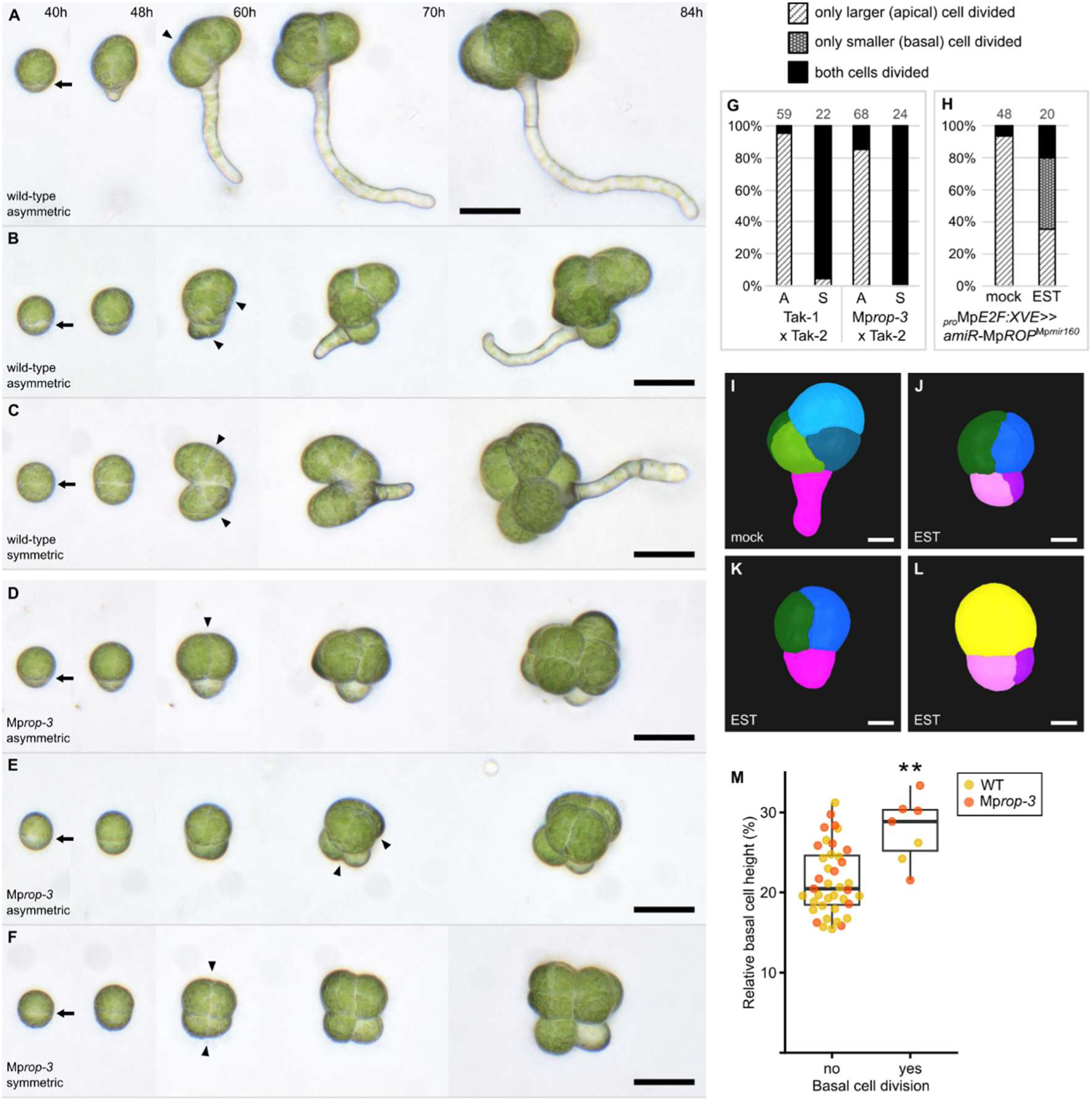
Highly asymmetric spore division, ensured by MpROP, promotes terminal cell differentiation. **(A–F)** Timelapse images of (A–C) wild type from the Tak-1 x Tak-2 population, and (D–F) Mp*rop-3* from the Mp*rop-3* x Tak-2 population. Arrows mark the spore division plane. Arrowheads mark the division planes of the second round of division. **(G)** Frequencies of only one or both daughter cells dividing following asymmetric (A) or symmetric (S) spore divisions in Tak-1 x Tak-2 and Mp*rop-3* x Tak-2 populations. Daughter cell divisions were assessed through timelapse imaging between 40–84 h after spore plating (A–F). Numbers above each bar indicate sample sizes. **(H)** Frequencies of one or both daughter cells dividing in the isogenic *_pro_*Mp*E2F:XVE*>>*amiR*-Mp*rop*^Mp*mir160*^ population treated with DMSO (mock) or estradiol (EST). Daughter cell divisions were assessed through confocal imaging of 72-h-old sporelings, which were fixed, cleared and stained with the cell wall dye SR2200 (I–L). **(I–L)** 3D segmented images of 72-h-old *_pro_*Mp*E2F:XVE*>>*amiR*-Mp*rop*^Mp*mir160*^ sporelings that were treated with DMSO (I) or EST (J–L). Both daughter cells divided (J); only larger daughter cell divided (K); only smaller daughter cell divided (L). **(M)** Relative basal cell height (proxy for division asymmetry) of asymmetrically divided spores from Mp*rop-3* x Tak-2, grouped based on if basal cell later divided. Relative basal cell height (basal cell height/total height) measured 40 h after spore plating. Basal cell division assessed through timelapse imaging between 40–84 h (D–E). ** indicates significant difference in the means of the two groups (Welch’s t-test: p = 5.0 x 10^-3^). Data points for individuals predicted to be wild type and Mp*rop-3* are colour coded in yellow and orange, respectively. n = 39 for non-divided and 7 for divided. Scale bars, 50 µm in (A–F); 10 µm in (I–L).

If a high degree of spore division asymmetry suppresses basal cell division and promotes its terminal differentiation, we hypothesised that basal cell division would occur more frequently in Mp*rop* mutants, where asymmetric spore divisions are on average more symmetric than in wild type (Figures 2E and 2F). To test this hypothesis, we tracked the fates of asymmetrically divided spores from the Mp*rop-3* x Tak-2 population, up to 84 h after spore plating. Basal cell division was observed in 15% (n = 10/68) of cases, significantly higher than the 5% (n = 3/59) observed in the pure wild-type Tak-1 x Tak-2 population (Figures 4B, 4E and 4G). To independently test if compromised MpROP function increases the frequency of basal cell division, we fixed 72-h-old *_pro_*Mp*E2F:XVE*>>*amiR*-Mp*rop*^Mp*mir160*^ sporelings, that were grown either in the presence or absence of estradiol. In the absence of estradiol, both daughter cells had divided in 6% (n = 3/48) of sporelings (Figure 4H). By contrast, in the presence of estradiol, both daughter cells had divided in 20% (n = 4/20) of sporelings (Figures 4H and 4J). Furthermore, exclusively in the estradiol treated population, we observed sporelings where the smaller, but not the larger, daughter cell had divided (Figure 4L). We interpret these as cases where a relatively symmetric spore division was followed by the division of the basal cell, and the sporelings were fixed before the eventual division of the apical cell. Taken together, these data support the hypothesis that MpROP suppresses basal cell division by promoting highly asymmetric spore division.

To determine if basal cell division in the *Mprop-3* x Tak-2 population is associated with reduced spore division asymmetry, we compared the degree of asymmetry of spore divisions that formed basal cells that either subsequently divided or terminally differentiated as rhizoid cells. From the time-lapse images of asymmetrically divided spores used to track daughter cell fates (Figures 4D and 4E), we measured spore division asymmetry as a ratio of basal cell height to total height (relative basal cell height). Asymmetric spore divisions that produce a dividing basal cell tended to be more symmetric than spore divisions that produce a terminally differentiating basal cell (Figure 4M). Furthermore, individuals predicted to be Mp*rop-3*, based on their distinctive isotropic growth phenotype, were overrepresented in the group with a divided basal cell. We conclude that MpROP promotes highly asymmetric spore divisions, and that this high degree of asymmetry promotes terminal differentiation of the basal cell.

### The two daughters of the apical cell differ in growth trajectories and MpROP localisation patterns

In contrast to the terminal differentiating basal cell, the larger apical cell divides to produce two apical daughter cells (Figure 1B). To determine if the fates of these two apical daughter cells differ with respect to their cell division potential, we examined the division patterns of 72-h-old wild-type sporelings comprising five or more cells. Out of 22 sporelings, we inferred both apical daughter cells had divided in 21 cases (Figures 5A and S5). This suggests that both apical daughter cells are specified to divide. To test if the two apical daughter cells differ in growth form, we tracked their growth through time-lapse imaging of 3-celled sporelings expressing Venus-MpROP, mScarletI-AtLTI6b (plasma membrane marker) and mScarletI-N7 (nuclear marker). Following apical cell division, we consistently detected the polarised outgrowth of just one daughter cell (Figure 5B; n = 22/29). Venus-MpROP was enriched at the cortical site of growth in this cell, indicating that MpROP localisation is correlated with localised, polarised growth (Figure 5B’). By contrast, the other daughter cell grew less and in an isotropic manner. Venus-MpROP polarisation was not detected at the cell cortex in this apical daughter cell. The shape and organisation of cells in the 5-celled wild-type sporeling also indicate that one apical daughter cell grew anisotropically before dividing whilst the other grew isotropically before dividing. (Figure 5A). We therefore conclude that the two apical daughter cells grow differently – one undergoing anisotropic growth in which a MpROP crescent forms and the other undergoing isotropic growth.

**Figure 5.**
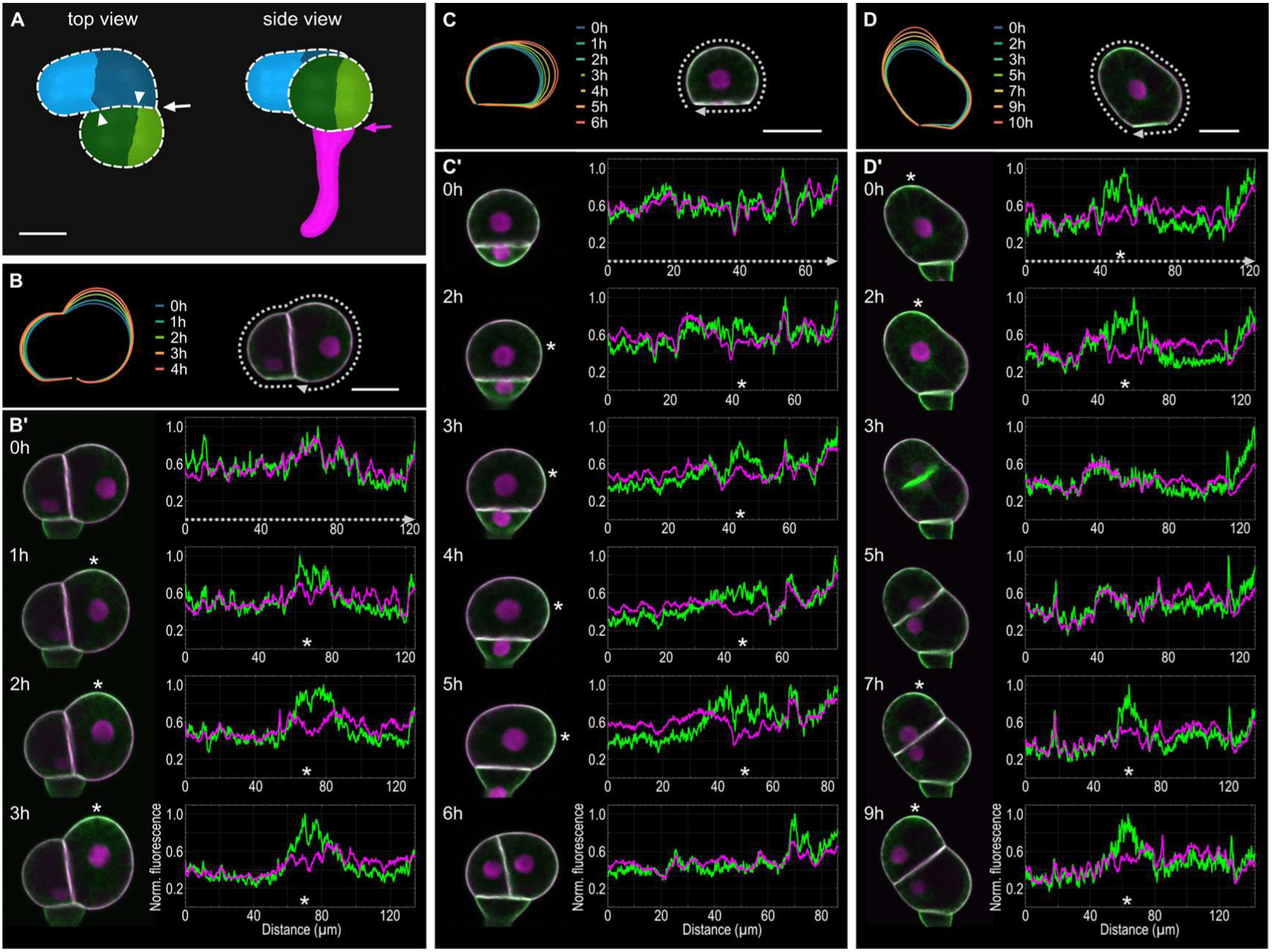
Polar localisation and asymmetric inheritance of Venus-MpROP in apical daughter cells. **(A)** 3D cell segmentation of representative 5-celled wild-type sporeling (4 cells in early cell mass). Division pattern inferred from cellular arrangement indicates both apical daughter cells divided. White dashed lines outline the two clonal halves of the early cell mass, each derived from one apical daughter cell. Pink arrow: first spore division; white arrow: apical cell division; white arrowheads: divisions of apical daughter cells. As the walls marked by the white arrowheads bisects the wall marked by the white arrow, they must have formed after the apical cell division. **(B–D)** Left: outlines of apical cell(s) in B’–D’ undergoing polarised outgrowth. Right: Grey dotted lines indicate the cell surface along which Venus and mScarletI fluorescence was qantified in B’–D’. **(B’–D’)** Left: Time-lapse images of sporelings expressing *_pro_*Mp*ROP:Venus-*Mp*ROP*, *_pro_*Mp*UBE2:mScarletI-*At*LTI6b*, and *_pro_*Mp*ROP:mScarletI-N7*. Right: Normalised fluorescence intensity profiles of Venus-MpROP (green) and mScarletI-AtLTI6b (magenta) along the cell surface marked by the grey dotted lines in B–D. Asterisks mark site of Venus-MpROP polarisation. All scale bars, 20 µm. See also Figure S5.

To test if the anisotropically growing apical daughter cell inherits the MpROP polarity crescent from the apical cell, we tracked Venus-MpROP localisation from before to after apical cell division. Early in the two-celled stage, when basal cell tip growth initiates, Venus-MpROP was localised uniformly at the apical cell cortex (Figure 5C’, 0h). Venus-MpROP then became enriched to the cortical region of the apical cell that initiated polarised outgrowth (Figures 5C’, 5D’). This demonstrates that local enrichment of MpROP is correlated with localised cell growth of the apical cell. Later, during cytokinesis, Venus-MpROP polarisation was lost at the cell cortex and Venus-MpROP marked the expanding cell plate, which formed perpendicular to the apical cell growth axis (Figures 5C’, D’). Following cytokinesis, the Venus-MpROP polarity crescent reestablished at or near the cortical region that harboured the Venus-MpROP polarity crescent in the apical cell, and polarised outgrowth reinitiated (Figure 5D’). The other daughter cell, which inherited the cortical region that was not previously enriched in Venus-MpROP, did not form a Venus-MpROP polarity crescent and grew isotropically. These data suggest that the contrasting growth trajectories of the two apical daughter cells are specified by the cortical regions inherited from the apical cell, with the daughter cell inheriting the region previously enriched in Venus-MpROP specified for anisotropic growth.

### MpROP-mediated anisotropic growth is required for the formation of two morphologically distinct halves in the early cell mass

The correlation between Venus-MpROP polarisation and localised growth suggested that MpROP promotes anisotropic growth in one of the two apical daughter cells. We therefore hypothesised that in the absence of MpROP function, both daughter cells would grow isotropically, resulting in the loss of morphological asymmetry between the two clonal halves of the early cell mass, which are each derived from one apical daughter cell. To test this, we segmented confocal images of 19 wild-type and 15 Mp*rop-3* sporelings comprising 4–8 cells in the early cell mass (Figures 6A and 6B). Based on the pattern of cellular arrangement, we inferred cells derived from the same apical daughter cell and grouped them to reconstruct the two clonal halves in 3D (Figure S5). We then quantified the morphological anisotropy of each clonal half (Figures 6C and 6D) and calculated the relative difference between each half as an asymmetry index. The asymmetry index was significantly greater for wild-type sporelings compared to Mp*rop-3* sporelings (Figure 6E). This is consistent with the hypothesis that MpROP function is required for the different growth forms of the two apical daughter cells and their clonal derivatives.

**Figure 6.**
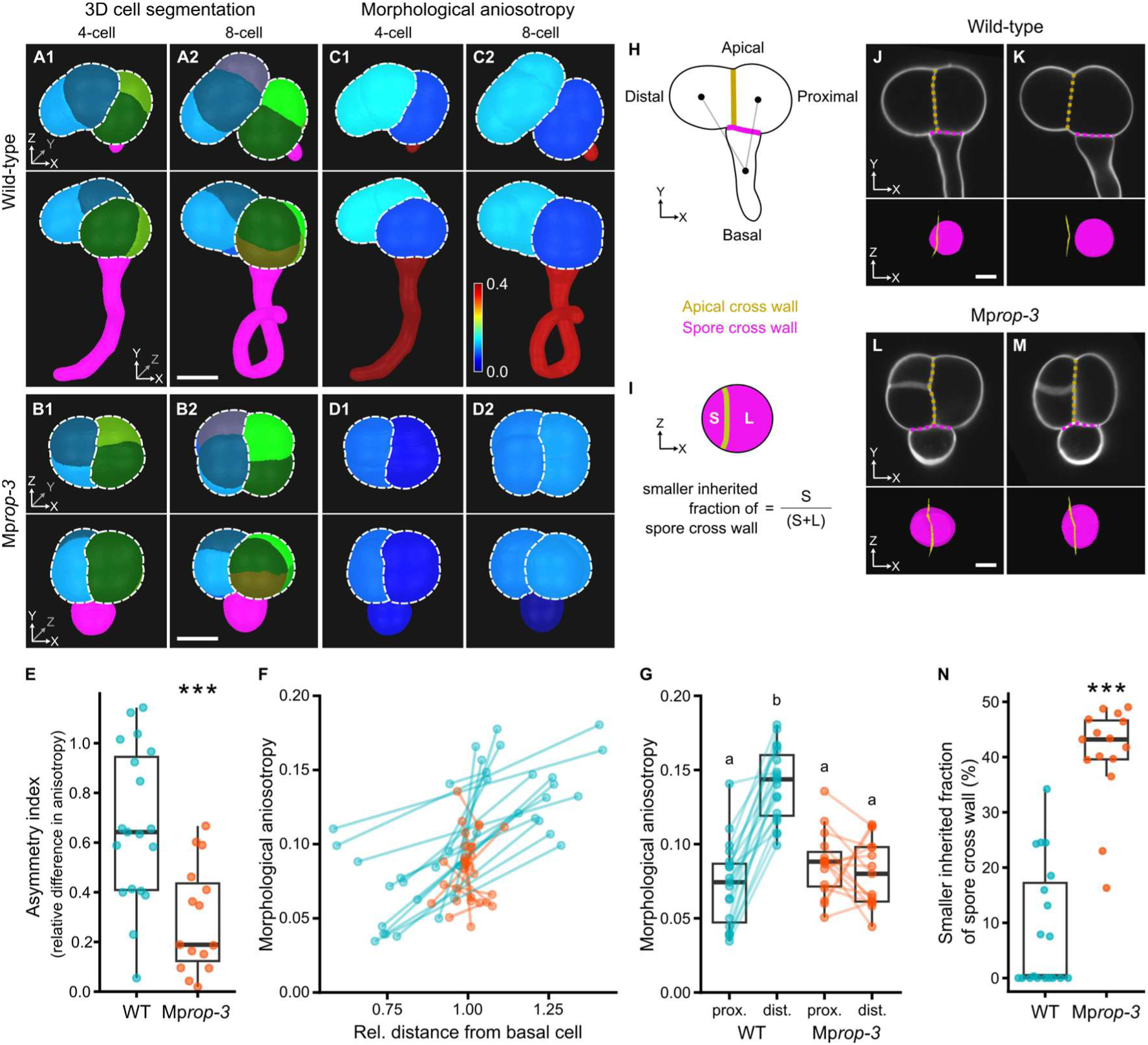
MpROP-mediated polarised outgrowth generates asymmetry within the early cell mass. **(A–B)** 3D cell segmentation of wild-type and Mp*rop-3* sporelings with 4 or 8 cells in the ealy cell mass. Upper and lower pannels show top and side views, respectively. White dashed lines outline the two clonal halves of the early cell mass. **(C–D)** Heatmap represenation of morphological anisotropy for the two clonal halves of sporelings in A–B. **(E)** Assymmetry index (defined as the relative differnece in morphological anisotropy between the two clonal halves) is significantly greater in wild type than in Mp*rop-3* (Student’s two-sample t-test: p = 3.6 x 10^-4^). Each dot represents measurement for one sporeling. **(F)** Morphological anisotropy for the two clonal halves plotted against their relative distance from the basal cell centroid. Two dots connected by a line represent the two clonal halves from the same sporeling. In wild type (blue), the more anisotropic clonal half is further from the basal cell centroid. **(G)** Morphological anisotropy for the proximal and distal halves of the early cell mass. See (H) for definitions of proximal and distal. Two dots connected by a line represent the two clonal halves from the same sporeling. Different letters indicate significantly different means (two-way repeated measures ANOVA followed by Tukey’s HSD test: p < 0.05). **(H)** Schematic representation of the proximal-distal asymemtry in the early cell mass. Dots indicate the ceontroids of the basal cell and the two apical halves. The distal half is defined as the apical half with a centroid further from the basal cell centroild. **(I)** Schematic representation of the spore cross wall surface (pink) bisected by the apical cross wall (yellow). S and L denote the smaller and larger fractions of the spore cross wall partitioned by the apical cross wall. Partitioning asymmetry was quantified as the smaller inherited fraction relative to the total spore cross wall area (see N). **(J–M)** Upper pannels show optical cross section images of 5-celled sporelings perpendicular to the spore cross wall surface (pink dotted line). Lower pannels show surface views of the spore cross wall (pink) bisected by the apical cross wall (yellow line). **(N)** The spore cross wall is partitioned asymmetrically between the two apical daughter cells in wild type but symmetrically in Mp*rop-3* (Welch’s t-test: p = 3.8 x 10^-8^; Wilcoxon rank-sum test: p = 3.8 x 10 ^-6^). Values were caluclated with the formula in (I). A value of 0 indicates the apical cross wall did not bisect the spore cross wall (as in K). n = 19 for wild type (Tak-1 × Tak-2) and 15 for Mp*rop-3* (Mp*rop-3* × Tak-2) in (E–G) and (N). Scale bars, 20 µm in (A–D); 10 µm in (J–M). ***p < 0.001.

To determine how the contrasting growth forms of the two apical daughter cells contribute to shaping the early sporeling body plan, we compared the positions of the two clonal halves relative to the basal cell. In wild-type sporelings, the centroid of the more anisotropic apical half was consistently further from the basal cell centroid than that of the less anisotropic apical half (Figures 6F and 6H). This establishes asymmetry within the early cell mass – a distal half, further from, and a proximal half, closer to, the basal cell centroid (Figure 6H). Unlike in wild type, the distances from the centroids of the two apical halves to the basal cell centroid were similar in Mp*rop-3* sporelings (Figure 6F). Moreover, the two halves had indistinguishable morphological anisotropy, both resembling the proximal half of wild type (Figure 6G), suggesting that in the absence of MpROP function, both apical daughter cells and their derivatives undergo isotropic growth. This indicates that by promoting polarised outgrowth of one apical daughter cell and its derivatives, MpROP contributes to the establishment of proximal-distal asymmetry within the early cell mass.

### MpROP contributes to establishing proximal-distal asymmetry in the apical cell

To determine if the proximal-distal asymmetry within the early cell mass is already established when the apical cell divides, we characterised the geometry of wild-type apical cell division. We found no evidence that the apical cell division is as highly asymmetric as the spore division, however, the relative position of the cross wall produced from the division of the apical cell (the apical cross wall) varied. It either attached to the first cross wall (the spore cross wall), or attached to the wall that forms the external surface of the apical cell (Figures 5B’–5D’, and 6H). In the former case, both apical daughter cells inherit portions of the spore cross wall, and in the latter, one daughter inherits the entire spore cross wall. We quantified how the spore cross wall area was partitioned between the two apical daughter cells in 20 wild-type sporelings (Figure 6I). In 9 cases, the apical cross wall bisected the spore cross wall (Figure 6J). In 8 of these 9 cases, the apical cross wall bisected the spore cross wall asymmetrically, with the distal apical half inheriting less than 25% of the spore cross wall area (Figure 6N). In the remaining 11 cases, the apical cross wall did not bisect the spore cross wall (Figure 6K) – the entire spore cross wall area was inherited by the proximal apical half. These data demonstrate that apical daughter cells form quantitatively different cell contacts with the basal cell, with the proximal half consistently sharing greater cell wall area with the basal cell. Moreover, the asymmetric placement of the apical cross wall with respect to the spore cross wall indicates that the proximal-distal asymmetry within the early cell mass is established by the time the apical cell divides.

Since Venus-MpROP is enriched at the cortical site of apical cell outgrowth, and the apical cross wall is positioned perpendicular to the apical cell growth axis (Figure 5), we hypothesised that the asymmetric placement of the apical cross wall relative to the spore cross wall, is dependent on MpROP function. To test this, we quantified how the spore cross wall area is partitioned between the two apical daughter cells in the Mp*rop-3* mutant. In 15 out of 15 cases, the apical cross wall bisected the spore cross wall (Figures 6L–6N). In 13 out of these 15 cases, the apical cross wall bisected the spore cross wall roughly symmetrically (with each daughter cell inheriting over 35% of the spore cross wall area). This contrasts with the cases in wild type, where the majority of apical cross walls either bisected the spore cross wall asymmetrically, or did not bisect at all. This demonstrates that MpROP is required for the asymmetric placement of the apical cross wall relative to the spore cross wall, and therefore that MpROP is involved in establishing the proximal-distal asymmetry in the apical cell, presumably by promoting polarised outgrowth.

Taken together, we conclude that MpROP contributes to initially establishing the proximal-distal asymmetry within the apical cell, leading to the formation of proximal and distal apical daughter cells with unequal contact with the basal cell. Subsequently, the two daughter cells develop unequal growth trajectories, with the distal cell undergoing MpROP-mediated polarised growth. This ultimately generates morphological asymmetry within the early sporeling body plan.

## DISCUSSION

We discovered that the MpROP protein polarises to the basal pole of the Marchantia spore before mitosis, where it contributes to ensuring the asymmetry of spore division. This physical asymmetry is necessary for the first cell differentiation event in the development of the multicellular Marchantia gametophyte. Furthermore, we show that MpROP-mediated polarised outgrowth generates asymmetry within the early cell mass that later gives rise to the prothallus. Our work represents one of the first genetic studies into how asymmetry is generated during early development of a multicellular plant body from a single isolated cell.

### Asymmetric spore division serves to produce a sufficiently small basal cell for terminal differentiation

By comparing daughter cell fates following asymmetric or symmetric spore divisions, we demonstrate that the physical asymmetry of Marchantia spore division is necessary to specify the terminal differentiation of the smaller basal cell into the first rhizoid cell (Figure 4). This resembles the case in the fern *Onoclea sensibilis*, where an asymmetric spore division generates a small terminally differentiating rhizoid cell, whereas an artificially induced symmetric spore division produces two daughter cells that continue to divide^25^. Asymmetric cell divisions where the smaller daughter cell is committed to differentiate and the larger daughter retains proliferative capacity are also observed in other diverse organisms, including *Drosophila melanogaster* and *Volvox carteri*^26,27^. This suggests the generation of a small cell and a large cell through asymmetric division represents a fundamental strategy for producing differentiating and proliferative cell types.

Utilising Mp*rop* mutants, where asymmetric spore divisions are on average quantitatively more symmetric than in wild type (Figure 2), we demonstrate a quantitative link between the extent of division asymmetry and cell fate (Figure 4). The increased frequency of basal cell division in Mp*rop* mutants compared to wild type indicates that a sufficiently high degree of spore division asymmetry – in other words, a sufficiently small basal cell volume relative to the apical cell volume – is required to reliably suppress basal cell division and promote terminal differentiation. This is consistent with the presence of a relative cell-size threshold, which the basal cell must fall below, for terminal differentiation to be triggered. In the *A. thaliana* stomatal lineage, a meristemoid formed through an asymmetric cell division differentiates into a guard mother cell if its volume is lower than a critical size threshold, but continues to divide if above the size threshold^28^. It has been suggested that terminal differentiation is triggered when a nuclear differentiation factor becomes sufficiently concentrated in the meristemoid as cell size falls below the critical threshold. Given that the highly asymmetric division of the Marchantia spore generates a higher nuclear to cell volume ratio in the basal cell compared to the apical cell (Figure 3D), it is plausible that this serves to achieve a sufficiently high (absolute or relative to apical cell) concentration of a nuclear differentiation factor in the basal cell.

### ROP likely reinforces spore polarity ensuring robust asymmetric spore division

We discovered that MpROP transiently accumulates at the spore basal pole just before mitosis, when the nucleus is basally positioned (Figures 1E and 1F). It remains unclear how or when spore polarity is first established, but the timing of MpROP polarisation – shortly before mitosis – coincides with when the apical-basal polarity axis becomes fixed^11^. The relatively mild defects in asymmetric spore division in Mp*rop* mutants also indicate that ROP is not required for initially establishing spore polarity (Figure 2). Rather, our data suggest that ROP contributes to reinforcing or maintaining cell polarity, potentially by securing the nucleus at the basal pole. This function is likely mediated through actin. ROP and actin work together to position the nucleus before asymmetric cell division in moss^29^, and actin is also transiently polarised at the spore basal pole before mitosis^10^. Moreover, disrupting either actin or ROP function causes only a subset of spores to divide symmetrically, indicating that neither is strictly required for polarity establishment but both contribute to the robustness of asymmetric division^10^.

Our finding contrasts with fungal and animal cells where RHO family GTPases play essential roles in polarity establishment. For example, in *S. cerevisiae*, CDC42 is required for cell polarisation and budding, which can be considered a form of asymmetric cell division^13,14^. CDC42 accumulates at a single cortical site through a positive feedback mechanism to define the site of bud emergence^15,16^. We propose that, rather than establishing spore polarity, ROP functions in a positive-feedback mechanism that reinforces cell polarity after its establishment, thereby ensuring that spore division is highly and reproducibly asymmetric.

### ROP-mediated polarised outgrowth generates asymmetry within the apical early cell mass

Following asymmetric spore division, MpROP polarises the growth of both the basal cell and the apical cell (Figures 3, 5, and 6). Similar to localised outgrowth promoted by animal and fungal RHO GTPases, this process is likely actin-mediated, which is consistent with the depolarisation of cell growth in sporelings treated with the actin depolymerising drug Latrunculin B^10^. Furthermore, we demonstrate that MpROP-mediated polarised outgrowth of the apical cell generates two daughter cells with unequal contact with the basal cell (Figure 6). The distal daughter, which retains little or no contact with the basal cell, undergoes MpROP-mediated anisotropic growth, whereas the proximal daughter, which retains greater contact with the basal cell, undergoes isotropic growth. Consequently, MpROP contributes to the formation of proximal-distal asymmetry within the apical early cell mass.

Our analysis of sporelings up to 3 days old indicate that the distal apical daughter gives rise to the sporeling apex, the region furthest from the germ rhizoid (Figure 6). In 4–5-day-old sporelings, the flat plant body – the prothallus – initiates formation at the sporeling apex with an auxin minimum^8^. Although we cannot rule out the possibility that the proximal apical daughter cell later gives rise to a new sporeling apex, the distal polarisation of Venus-MpROP promoting polarised outgrowth in the apical cell is inherited by the distal daughter, suggesting that the proximal-distal polarity set up in the apical cell is maintained after division (Figure 5D). We therefore propose that the maintenance of this polarity axis biases prothallus initiation to the distal end of the early cell mass.

How the unequal contacts of the two apical daughter cells with the germ rhizoid influence their fates and the subsequent pattern of sporeling morphogenesis remains unclear. However, it is consistent with a model in which intercellular signalling, via mobile signals, takes place between the early cell mass and the germ rhizoid. The unequal contact could contribute to establishing asymmetric distribution of mobile signals within the early cell mass. This in turn could make different parts of the early cell mass develop differently. Thus, the unequal cell contacts generated as a consequence of MpROP-mediated polarised growth, may provide a means to generate further asymmetry to pattern the early sporeling.

## ACKNOWLEDGMENTS

We thank the GMI/IMBA/IMP BioOptics team for advice on confocal microscopy and image analysis and for assistance on FACS. We thank Molecular Biology Services, the Media Lab, and the Lab Support of GMI/IMBA/IMP, and the VBCF Plant Sciences unit for their support. We thank Eri Okada (Kobe University) for technical assistance. This work was funded by a grant in aid from the Austrian Academy of Sciences (OEAW) to the Gregor Mendel Institute and by a European Research Council Advanced Grant (DENOVO-P, project no. 787613) to L.D. H.M. was supported by a Biotechnology and Biological Sciences Research Council Doctoral Training Partnership Scholarship (grant no. BB/M011224/1) and an EMBO Postdoctoral Fellowship (ALTF 490-2024). Z.M. was funded by a Human Frontier Science Program Fellowship (LT0024-2023) and an EMBO Postdoctoral Fellowship (ALTF 839-2022). Y.S. was supported by JSPS KAKENHI grants (21J40092, 22KJ2244, and 25K09689). Y.S. and K.I. were supported by GteX Program Japan (JPMJGX23B0).

## AUTHOR CONTRIBUTIONS

H.M. and L.D. designed the project. Y.S. and K.I. designed the experiments using the conditional Mp*ROP* knock-down line. H.M. and Y.S. generated transgenic spores, performed the experiments, and conducted the formal analyses. K.J. assisted H.M. with plant propagation, spore generation, genotyping, and image analysis. Z.M. and M.M. established the FACS spore plating method. H.M. wrote the original draft with input from L.D. All authors contributed to reviewing and revising the manuscript.

## MATERIALS AND METHODS

### Plant lines used and crossing schemes

A wild-type population of spores was generated by crossing the Takaragaike-1 (Tak-1, male) and Takaragaike-2 (Tak-2, female) accessions of *Marchantia polymorpha*^30^. All transgenic spores used in this study were likewise in a mixed Tak-1/Tak-2 genetic background. All parental lines, excluding *_pro_*Mp*E2F:XVE*>>*amiR*-Mp*rop*^Mp*mir160*^, were previously generated^10,22^. A segregating population of Mp*rop-3* and wild-type spores was generated by crossing a Cas9-free male Mp*rop-3* plant^22^ with Tak-2 (Figure S1A). An isogenic population of spores expressing *_pro_*Mp*E2F:XVE*>>*amiR*-Mp*rop*^Mp*mir160*^ was generated by first crossing a male T_0 *pro*_Mp*E2F:XVE*>>*amiR*-Mp*rop*^Mp*mir160*^ plant with Tak-2 to obtain a female T_1 *pro*_Mp*E2F:XVE*>>*amiR*-Mp*rop*^Mp*mir160*^ plant, which was then backcrossed with the T_0_ plant (Figure S2D). Spores expressing *_pro_*Mp*ROP:Venus-*Mp*ROP*, *_pro_*Mp*UBE2:mScarletI-*At*LTI6b*, and *_pro_*Mp*ROP:mScarletI-N7* were generated by crossing a male *_pro_*Mp*ROP:Venus-*Mp*ROP*^22^ with a female line carrying a single insertion encoding both *_pro_*Mp*UBE2:mScarletI-*At*LTI6b* and *_pro_*Mp*ROP:mScarletI-N7*^10^.

### Plant growth conditions

Plants were vegetatively propagated through gemmae or thallus clippings on ½-strength B5 Gamborg’s medium (1.5 g/L B5 Gamborg, 0.5 g/L MES hydrate, 1% sucrose, pH adjusted to 5.5, solidified with 1% plant agar) under sterile conditions. Plants on agar plates were grown at 22– 23 ℃ under continuous white light (50–60 µmol m^-2^ s^-1^).

For spore production, gametangiophore formation was induced by far-red light irradiation based on the protocol described by Chiyoda et al.^31^ To generate *_pro_*Mp*E2F:XVE*>>*amiR*-Mp*rop*^Mp*mir160*^ spores, two-week-old gemmalings were transplanted onto vermiculite wetted with a 1:2,000 dilution of Hyponex fertilizer in plastic containers (Risupack). A ventilation opening was made in the lid and covered with surgical tape. Plants were grown for a further week under continuous white light and then transferred to continuous white light (60–70 µmol m^-2^ s^-1^) supplemented with far-red light (20–30 µmol m^-2^ s^-1^) at 20 ℃.

To induce gametangiophore formation in all other lines, plants were grown at 20 ℃ and 60% humidity under long-day conditions (16 h light, 8 h dark), with white light (50–60 µmol m^-2^ s^-1^) supplemented with far-red light (30–40 µmol m^-2^ s^-1^). Mp*rop-3* plants were grown on sterile ½-strength B5 Gamborg’s medium in MagentaTM GA-7 boxes (Sigma) and watered regularly to prevent the plants from drying out. Two- to three-week-old gemmalings of the remaining lines were transplanted onto autoclaved soil (1:3 mixture of fine vermiculite and Neuhaus N3 compost) in SacO2 Microbox containers as previously described^22^.

Spores were grown on a sterilised sheet of cellophane placed on the surface of ½-strength B5 Gamborg’s medium at 22–23 ℃ under continuous white light (50–60 µmol m^-2^ s^-1^). To induce conditional expression of amiR-MpROP ^Mp*mir160*^, *_pro_*Mp*E2F:XVE*>>*amiR*-Mp*rop*^Mp*mir160*^ spores were cultured on a sterilised sheet of cellophane placed on the surface of ½-strength B5 medium containing 10 μM β-estradiol (Fujifilm) or the equivalent volume [0.1% (v/v)] of DMSO (Fujifilm) as a mock control.

### Generating conditional Mp*ROP* knockdown line

To generate the *_pro_*Mp*E2F:XVE*>>*amiR*-Mp*rop*^Mp*mir160*^ transgenic line, an artificial microRNA targeting the Mp*ROP* transcript was designed using the Mp*MIR160* backbone with the amiRNA Design Helper tool available on the MarpolBase website^32,33^(Figures S2A–S2C). An annealed pair of synthesized 85-mer oligonucleotides for the sequence spanning from the miR to the miR* (Oligo_F_amiR_MpROP and Oligo_R_amiR_MpROP) of *amiR*-Mp*rop*^Mp*mir160*^ was cloned into the PaqCI cloning site of the pMpAmiR_160_En01 vector, which contains the Mp*MIR160* backbone flanked by Gateway attL1 and attL2 sites^23^. The insert flanked by attL1 and attL2 sites was transferred into the binary vector pMpGWB168 by Gateway LR reaction^34^. The resulting vector encoding *_pro_*Mp*E2F:XVE*>>*amiR*-Mp*rop*^Mp*mir160*^ was used to transform regenerating Tak-1 thalli as previously described^35^.

### Harvesting sporangia and sterilising spores

Sporangia were harvested in 1.5-ml Eppendorf tubes, dried with silica gel, then stored at −70 to −80 °C until use as previously described^9,22^. *_pro_*Mp*E2F:XVE*>>*amiR*-Mp*rop*^Mp*mir160*^ spores were sterilised with 0.2% sodium hypochlorite (NaClO) supplemented with 0.05% Triton X-100 for 1 min, then pelleted by centrifugation (10,000 x g, 1 min). Spores of all other lines were sterilised with 0.1% sodium dichloroisocyanurate (NaDCC) for 2 min, then pelleted by centrifugation (15,000 x g, 2 min). After centrifugation, the sterilisation solution was removed and the spores were resuspended in sterile water.

### Plating spores using FACS

After sterilisation, spores were filtered through a 50 µm cell strainer (Sysmex) and resuspended in 1 ml sterile water. Spore solution was taken for sorting and plating of individual spores on an agar medium plate by BD FACS-Aria III Cell Sorter, using flow-cell nozzle of 70 µm diameter, and with the following sorting specifications: Cooling of both sample and collection plate were turned off, aiming to sort at room temperature. Sample Agitation set to 300rpm to ensure sample homogeneity throughout sorting. ND-filter x1.0 was used to allow proper detection of small spores, and a flow-rate of 5.0 was applied for sorting hundreds of individual spores within a few minutes. Two consecutive gates were applied to mark spores, FSC-A/SSC-A gate marking a distinct spore population based on size, and additional SSC-A/FSC-W gate excluding potential spore doublets or aggregates. Both gates included >90% of all events and resulted in the sorting of individual spores at high fidelity. Spores were distributed and spaced equally by the sorting layout on a sheet of sterilised cellophane placed on standard growth medium in PlusPlate (Singer Instruments). Following plating, the PlusPlate was sealed with its lid and left in standard growth conditions for seven days.

### Genotyping CRISPR mutants

To confirm the identity of Mp*rop-3* and wild-type progeny from the cross between Mp*rop-3* and Tak-2 (Figure S1F), genotyping was performed as previously described by Streubel et al.^36^ Primers sgRNA1_PCR_Fw and sgRNA1_PCR_Rv were used for PCR amplification and sgRNA1_Seq_Fw for Sanger sequencing^22^.

### Brightfield microscopy of live spores and sporeling

Brightfield images of live spores and sporelings were acquired with a Keyence VHX7000 digital microscope equipped with the VH-ZST lens and the VHX-7020 camera. For time-lapse imaging of individual spores grown on cellophane on ½-strength Gamborg’s medium, 3D depth composition images were acquired at x 1000 magnification, and the stage position for each spore was recorded at the first imaging timepoint. The media plates were returned to the growth chamber between imaging timepoints. To track cell fates following asymmetric or symmetric spore division, individuals where the spore cross wall was clearly visible and oriented perpendicular to the imaging plane were selected at the first time point. To image 7-day-old sporelings plated in a grid format using FACS, the 2D image stitching function was used at x 20 magnification.

### Sample fixation and clearing

To fix spores and sporelings, the piece of cellophane (2.5 cm x 2.5 cm) on which they were grown was peeled off the agar surface, rolled into a cylinder with the sample side facing inward, and placed in a 2-ml Eppendorf tube. The fixative (4% paraformaldehyde in 1x PBS supplemented with 0.1% Brij L23, or 4% formaldehyde (methanol free) in 1x PBS) was pipetted onto the inner cellophane surface to wash the samples off into the Eppendorf tube, after which the cellophane was removed. Following a 30 min incubation in 1 ml of the fixative, samples were pelleted through centrifugation (7,000 x g, 3 min) and the fixative was replaced with 1x PBS. After a few minutes in 1x PBS, the samples were pelleted (7,000 x g, 3 min), and the PBS was replaced with ClearSee-alpha^37^. Samples were left in the clearing solution at room temperature in the dark on a rocker for at least 5 days before imaging as previously described^38^.

### Confocal microscopy of fixed spores and sporelings

The day before imaging, Renaissance stain SR2200 was added to the samples in ClearSee-alpha to a final concentration of 0.2%. Before imaging, the samples were pelleted (7,000 x g, 3 min), most of the supernatant removed, and the samples were resuspended in 100 µl ClearSee-alpha. Resuspended *_pro_*Mp*E2F:XVE*>>*amiR*-Mp*rop*^Mp*mir160*^ spores were mounted using a 20 mm diameter Secure-Seal Spacer placed between two coverslips. For all other spores, resuspended samples were mounted using a 65 µl Gene Frame attached to a standard microscope slide.

Confocal imaging of Tak-1 x Tak-2 and Mp*rop-3* x Tak-2 progenies was performed on an inverted Zeiss LSM880 equipped with GaAsP detectors. Images were acquired with a LCI Plan-Apochromat 40×/1.2 NA objective using silicon immersion oil, which closely matches the refractive index of ClearSee-alpha^39^. SR2200 fluoresce was excited with the 405 nm laser, and emission in the range of 420–500 nm was collected. For 3D morphometric analysis in MorphoGraphX, 16-bit images were obtained with cubic voxels (0.4 x 0.4 x 0.4 µm).

Confocal imaging of *_pro_*Mp*E2F:XVE*>>*amiR*-Mp*rop*^Mp*mir160*^ samples was performed on a Leica STELLARIS 5. Images were acquired with a 20 × objective (HC PL APO CS2 20x/0.75 NA IMM) using silicon immersion oil. For 3D morphometric analysis in MorphoGraphX, images were obtained with voxel dimension of approximately 0.57 x 0.57 x 0.5 µm.

### Confocal microscopy of live spores and sporelings

Shortly before imaging, the piece of cellophane on which the spores were grown was peeled off the agar surface and transferred onto a thin agar slab (½-strength B5 Gamborg’s, 1% sucrose, 1% agar) placed within an imaging chamber described by Kirchhelle and Moore^40^. The chamber comprised a polydimethylsiloxane (PDMS) gum gasket stuck on a microscope slide, forming a well that was filled with liquid medium (½-strength B5 Gamborg’s, 1% sucrose).

Confocal live imaging of spores or sporelings expressing *_pro_*Mp*ROP:Venus-*Mp*ROP*, *_pro_*Mp*UBE2:mScarletI-*At*LTI6b*, and *_pro_*Mp*ROP:mScarletI-N7* was performed on an upright Zeiss LSM780 equipped with GaAsP detectors. For fluorescence quantification, 16-bit images were acquired with a Plan-Apochromat 20x/0.8 NA air objective. The following excitation laser wavelength and emission capture bandwidth were used: Venus (ex 514 nm, em 518–544 nm), mScarletI (ex 561 nm, em 571–624 nm). Sequential scanning was used to avoid bleed through. During time-lapse imaging, the imaging chamber was illuminated with cool white LED lights (approximately 45 µmol m^-2^ s^-1^) to promote plant development^7^.

### Segmentation and morphometric analysis in MorphoGraphX

For quantifying spore division asymmetry, images of only recently divided spores were analysed to minimise the influence of post-cytokinetic cell growth on the sizes of the apical and basal cells. Recently divided spores were identified based on a weaker SR2200 signal at the spore cross wall compared to the surrounding cell wall, a slightly undulating cross wall, and the absence of inward pinching of the parental cell wall at the junction with the spore cross wall.

3D cell segmentation was performed in MorphoGraphX 1.0^41^ or 2.0^42^ as follows. SR2200 fluorescence images were smoothed using a Gaussian blur (radius 0.3–0.5 µm), 3D segmented (ITK auto-seeded watershed), and finally a 3D mesh was generated with the Marching Cubes Algorithm (cube size: 0.8–1.0 µm, smooth pass: 3). For quantifying the shapes of the two clonal halves of the early cell mass, labels corresponding to cells within each clonal half were merged after watershed segmentation and before mesh creation.

Volume (*V*), Cell wall area (*A_c_*), Outside wall area (*A_o_*), Cell distance from the basal cell (*d*), and 3D shape anisotropy (*a*) were computed in MorphoGraphX 2.0 using the Mesh/Heat Map/Analysis/Cell Analysis 3D and Mesh/Cell Axis 3D/Shape Analysis/Compute Shape Analysis 3D processes. From these measurements, the following parameters were calculated. For recently divided spores: relative basal cell volume = *V*_basal_/(*V*_apical_ + *V*_basal_). For 72 h-old sporelings: basal cell sphericity = π^1/3^(6*V*_basal_)^2/3^/*A_c_*_, basal_; asymmetry index = |*a*_distal_ − *a*_proximal_|/((*a*_distal_ + *a*_proximal_)/2); relative distance from the basal cell = *d_i_*/((*d*_distal_ + *d*_proximal_)/2), where *i* is distal or proximal; wall area shared with neighbouring cells (*A_n_*) was calculated as *A_c_* – *A_o_*; wall area shared between basal cell and proximal half of early cell mass (*A_bp_*) = (*A_n_*_, basal_ + *A_n_*_, proximal_ – *A_n_*_, distal_)/2; wall area shared between basal cell and distal half of early cell mass (*A_bd_*) = (*A_n_*_, basal_ + *A_n_*_, distal_ – *A_n_*_, proximal_)/2; smaller inherited fraction of the spore cross wall = min(*A_bp_*, *A_bd_*)/(*A_bp_* + *A_bd_*).

### Morphometric analysis in Fiji

#### Rhizoid length measurement

Following 3D segmentation of 72 h-old fixed sporelings in MorphoGraphX, the segmented sporeling was spatially oriented so that the spore cross wall was perpendicular to the image plane and the whole length of the rhizoid was visible. The segmentation image was then imported into Fiji, where the segmented line tool was used to manually trace the rhizoid to measure its length.

#### Roundness and cross-sectional area of spores and sporelings

Time-lapse images of spores expressing *_pro_*Mp*ROP:Venus-*Mp*ROP*, *_pro_*Mp*UBE2:mScarletI-* At*LTI6b*, and *_pro_*Mp*ROP:mScarletI-N7* were first examined to identify spores with the apical-basal axis oriented parallel to the image plane and the spore cross wall perpendicular to it. For spores meeting these criteria, a sum projection image of the YFP channel Z-stack was generated and converted into a binary mask with holes filled. The Analyze Particles command in Fiji was then used to quantify the mask area and roundness.

### Fluorescence intensity quantification

To detect Venus-MpROP polarisation, fluorescence intensities of Venus-MpROP and mScarletI-AtLTI6b (a uniform plasma membrane marker) along the plasma membrane were measured and compared in Fiji as previously described^22^. For the analysis of single-timepoint spore images, we selected images where the nucleus was in the medial plane of the Z-stack. For time-lapse images of spores and sporelings, we selected those where the apical-basal axis was oriented parallel to the image plane and the spore cross wall perpendicular to it. A sum projection of five consecutive slices centred on the medial plane was generated for fluorescence measurements. The segmented line tool (line width: 2, spline fit) was used to manually trace the plasma membrane and fluorescence intensity profiles were generated using the Plot Profile command. To facilitate comparison of the Venus and mScarletI signal intensity profiles, the signal from each channel was background-subtracted, normalised so that the mean signal intensity was equal between channels, and finally scaled so that the maximum plotted intensity was 1.

## SUPPLEMENTAL FIGURES

**Figure S1.**
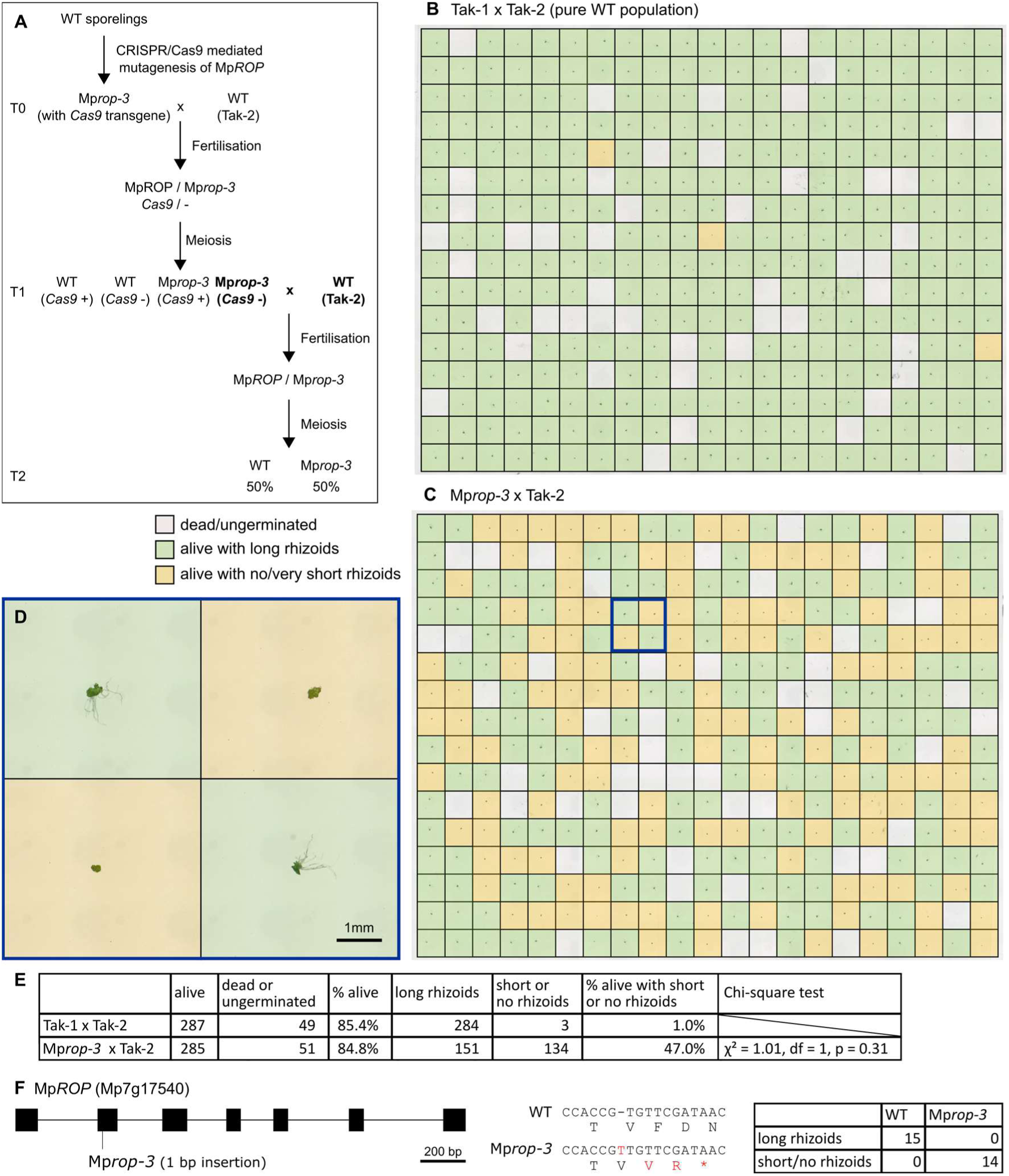
The cross between Mp*rop-3* and Tak-2 produces a 1:1 segregating population of viable Mp*rop-3* and wild-type progeny (related to Figure 2) **(A)** Crossing scheme for generating a 1:1 segregating population of Mp*rop-3* and wild-type spores. **(B–C)** Spores from Tak-1 x Tak-2 and Mp*rop-3* x Tak-2 crosses were plated on cellophane on solid media, in a 16 x 21 (336) grid format using FACS. Seven days after plating, sporelings were imaged to determine if they were dead or ungerminated (grey), alive with long rhizoids (green), or alive with no or very short rhizoids (yellow). **(D)** Zoomed in image of the region in C outlined with a blue box. **(E)** Summary of data from B and C. In the Mp*rop-3* x Tak-2 population, the observed segregation ratio of sporelings with and without long rhizoids (151:134) did not significantly differ from a 1:1 segregation ratio (chi-square test, p = 0.31). **(F)** From the Mp*rop-3* x Tak-2 cross, progeny with short or no rhizoids were all genotyped as Mp*rop-3* (n = 14) and progeny with long rhizoids were all genotyped as wild type (n = 15) through Sanger sequencing.

**Figure S2.**
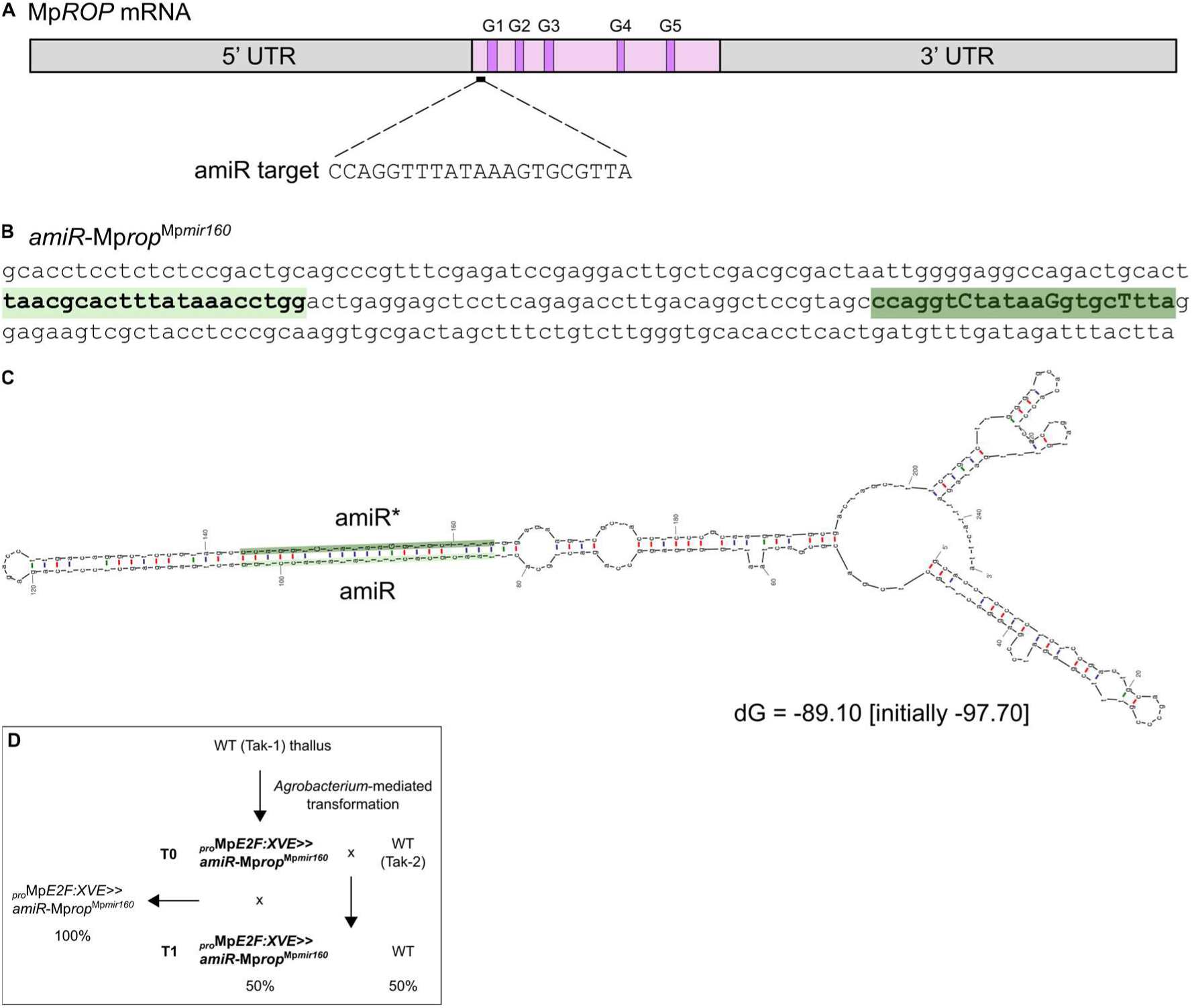
Generating an isogenic population of inducible Mp*ROP* knock-down spores (related to Figure 2) **(A)** Schematic of the Mp*ROP* (Mp7g17540) transcript showing the artificial microRNA (amiR) target site. The coding sequence is shown in pink, regions encoding the five highly conserved G boxes in purple, and the 5′ and 3′ untranslated regions (UTRs) in grey. **(B)** The amiR precursor sequence based on the Mp*MIR160* backbone (*amiR*-Mp*rop*^Mp*mir160*^). The amiR and amiR* sequences (light and dark green, respectively) were designed using the amiRNA Design Helper tool on MarpolBase (Tanizawa et al., 2026). **(C)** Predicted minimum free-energy (MFE) secondary structure of the amiR precursor. **(D)** Crossing scheme for generating an isogenic population of *_pro_*Mp*E2F:XVE>>amiR*-Mp*rop*^Mp*mir160*^ spores.

**Figure S3.**
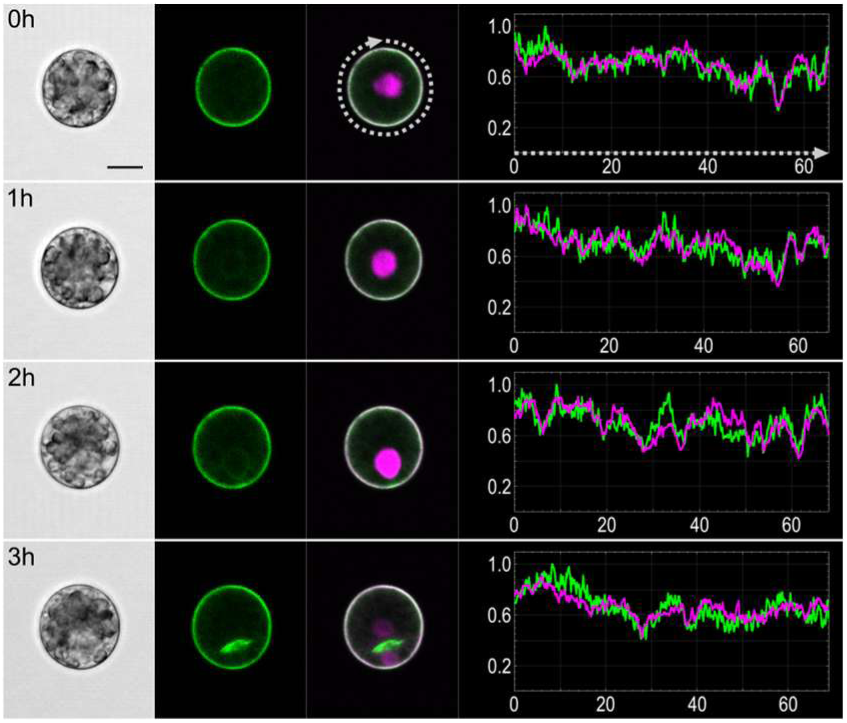
Venus-MpROP polarisation at the basal pole is lost or diminished during cytokinesis (related to Figure 3) Time-lapse images of a representative spore expressing *_pro_*Mp*ROP:Venus-*Mp*ROP*, *_pro_*Mp*UBE2:mScarletI-*At*LTI6b*, and *_pro_*Mp*ROP:mScarletI-N7*. From left to right: brightfield, Venus, and merged fluorescence images. Normalised fluorescence intensity profiles of Venus-MpROP (green) and mScarletI-AtLTI6b (magenta) along the spore perimeter (grey dotted line) are plotted for each time point. Normalised fluorescence intensity on the y-axis and distance (µm) on the x-axis. Scale bar, 10 µm.

**Figure S4.**
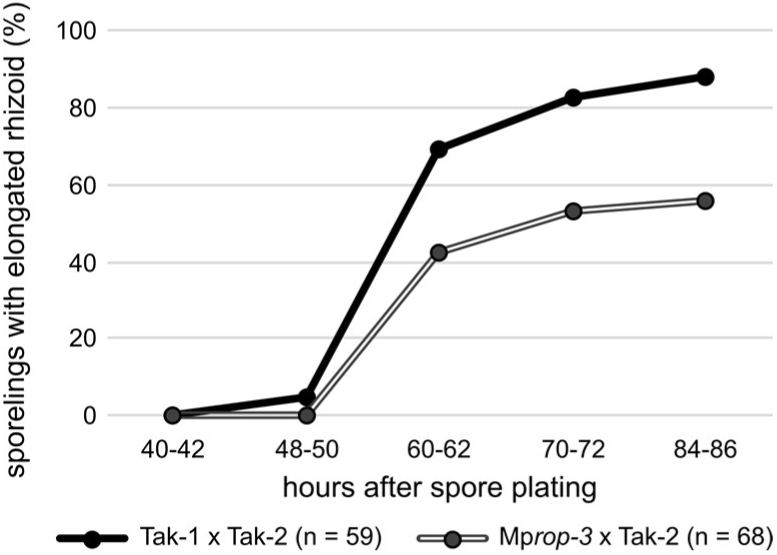
Sporelings with an elongated rhizoid following asymmetric spore division (related to Figure 3) Asymmetrically divided spores were identified 40–42 h after plating on cellophane on solid media. Basal cell development was tracked up to 84–86 h after spore plating to determine the proportion of sporelings that had formed an elongated rhizoid. A rhizoid cell was determined to be elongated if its length was at least twice its width.

**Figure S5.**
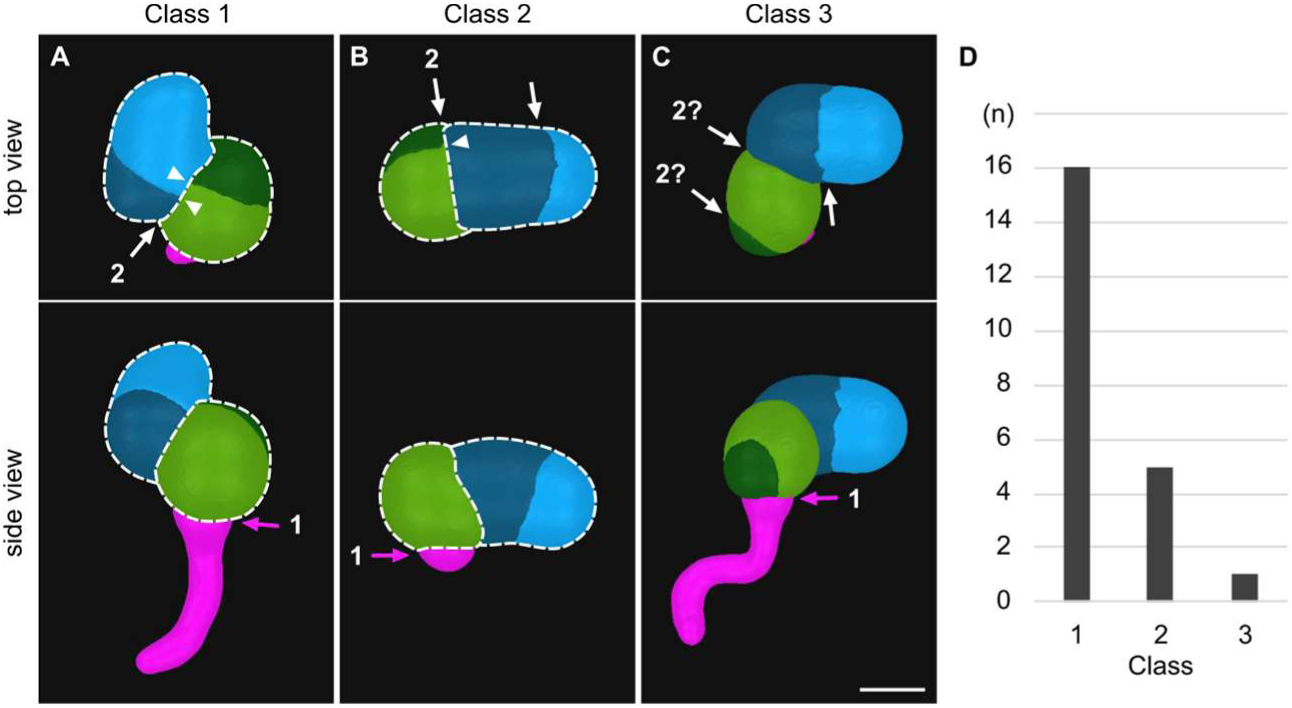
Inferring the order of cell divisions from 3D-segmented sporelings (related to Figure 5) 3D-segmented 72h-old wild-type sporelings with 4–8 cells in the early cell mass were categorised into three classes based on the pattern of cellular arrangement. Pink arrow marks the spore cross wall, inferred to have formed from the first division of the spore. The white arrow labelled 2 marks the cross wall inferred to have formed from the second division (apical cross wall). White dashed lines outline the two inferred clonal halves of the early cell mass in classes 1 and 2. Since a newly forming cross wall cannot bisect a cross wall that does not already exist, a cross wall that bisects another must have formed later. **(A)** Class 1: sporelings with one cross wall where its edge does not bisect the surface of another cross wall within the early cell mass (white arrow). All other cross walls (excluding the spore cross wall) bisect another cross wall within the early cell mass to form tricellular junctions (white arrowheads). Therefore, the white arrow marks the only geometrically possible apical cross wall. Its surface is bisected by at least one cross wall from either side, demonstrating that both apical daughter cells divided. **(B)** Class 2: sporelings with two cross walls which does not bisect the surface of another cross wall within the early cell mass (white arrows). During early sporeling development, division of the cell at the growing apex, rather than intercalary division is observed. Hence, the cross wall labelled 2 is inferred to have formed before the cross wall marked with the other white arrow and thus represents the apical cross wall. **(C)** Class 3: sporeling with no cross walls which bisect another cross wall within the early cell mass. The apical cross wall cannot be inferred with high confidence. Either cross wall labelled 2? could represent the third division without requiring an intercalary division. **(D)** Number of sporelings in each class (total n = 22). Scale bar, 20 µm.

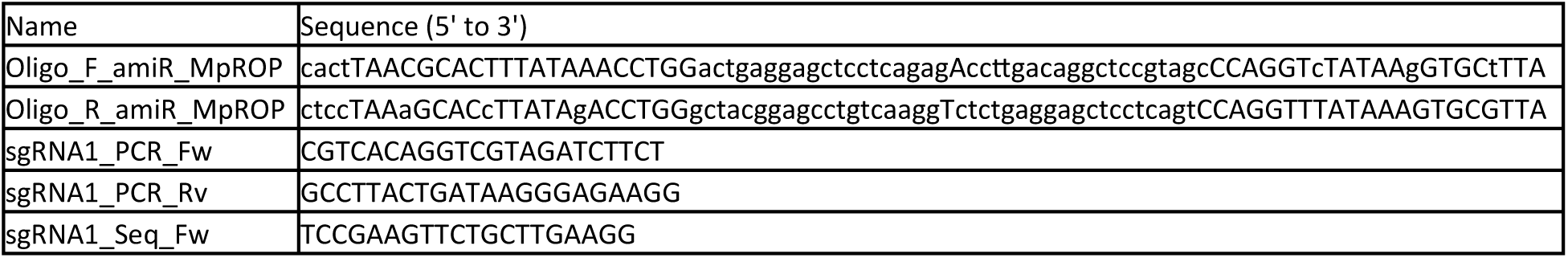

**Data S1. Sequences of oligonucleotides used in this study**

